# Blood, sweat, and beers: investigating mosquito biting preferences amidst noise and intoxication in a cross-sectional cohort study at a large music festival

**DOI:** 10.1101/2025.08.21.671470

**Authors:** Sara Lynn Blanken, Maartje R. Inklaar, Zhong Wan, Felix Evers, Merel Smit, Vladyslav Kalyuzhnyy, Julie M.J. Verhoef, Ezra T. Bekkering, Michelle Schinkel, Saskia Mulder, Carolina M. Andrade, Geert-Jan van Gemert, Alem Gusinac, Patrick Zeeuwen, Thomas H.A. Ederveen, Teun Bousema, Felix J.H. Hol

## Abstract

**Introduction:** We live in a world split between mosquito magnets and those lucky enough to remain (nearly) untouched. The reasons why some people attract more mosquito bites than others remain largely mysterious. In the Mosquito Magnet Trial, we investigated differences in mosquito attraction amongst festivalgoers with varying levels of hygiene and intoxication.

**Methods:** Our study was conducted in a slightly steamy pop-up laboratory inside four connected shipping containers at Lowlands Festival in The Netherlands (August 18–20, 2023). Participants completed an anonymous questionnaire on hygiene, diet, and festival-related behaviour (including alcohol uptake and shared sleeping arrangements). Mosquito attraction was measured using a custom designed setup: a transparent cage with perforations where female *Anopheles* mosquitoes were offered a choice between a sugar-feeder and the participants arm. Mosquitoes could only smell, not bite, the participant’s arm. Attraction was quantified through video imaging, measuring arm landings relative to total landings. Mosquito attraction was correlated with questionnaire responses and skin microbiota profiles collected from forearm skin swabs.

**Results:** Amongst the 465 included participants, mosquitoes showed a clear fondness for those who drank beer over those who abstained from the liquid gold (Fold Change 1.35, 95% CI 1.12–1.63, P_FDR_<0.001). Attraction was also contagious: participants that successfully lured a fellow human into their tent the previous night also proved more enticing to mosquitoes (FC 1.34, 95% CI 1.14–1.58, P_FDR_=0.002). Meanwhile, skipping the morning showering routine and using sunscreen reduced mosquito attraction (FC 0.52, 95% 0.38-0.70, P_FDR_<0.001). *Streptococci* were more abundant on the skin of highly attractive individuals (P_uncorrected_=0.017), and the overall abundance of malodour associated bacteria was high – possibly reflecting the container’s smell.

**Conclusion:** The Mosquito Magnet Trial, to our knowledge the largest study of its kind, was conducted in a loosely controlled setting with a selection bias towards science loving festivalgoers. That said, using our custom designed experimental set-up, we found that mosquitoes are drawn to those who avoid sunscreen, drink beer, and share their bed. They simply have a taste for the hedonists among us.

## INTRODUCTION

Do you get bitten by mosquitoes often, or are you lucky enough to escape their hunt for blood? The quest for factors behind observed differences in the preference of mosquitoes for certain humans has not only inspired scientists, it also forms a matter of public debate where factors as blood type, ‘sweet blood’, and alcohol consumption are regularly suggested. Mosquito nuisance was already described in the 5^th^ century BCE by the Greek historian Herodotus – long before their role in disease transmission was understood [1]. Today, it is well known that mosquitoes can carry pathogens which may be transmitted to human hosts during blood feeding. This makes mosquitoes more than just annoying in endemic regions; individuals who are bitten frequently face greater exposure to pathogens, increasing their risk of contracting mosquito-borne illnesses such as malaria or dengue.

Mosquitoes rely on their senses in order to find a human to bite. The search for a blood meal typically starts with the perception of CO_2_ that we exhale [2], which is considered the initial cue for mosquito activation. What then follows is a complex set of olfactory, visual, thermal, and physical cues that guide a mosquito to a human host [3, 4]. Sensory factors contributing to mosquito attraction can be universal, including CO_2_ and body heat, but also more specific to certain human individuals, like our smell. As most of us have experienced one way or the other, humans can have distinct odours. Odours depend on different skin microbiota and skin volatile compound compositions, which can influence mosquito attraction [5]. The skin microbiome of highly attractive people is reported to be of lower bacterial diversity [6] and producing higher amounts of volatile fatty acid metabolites [7] or long-chain carboxylic acids [8, 9]. In contrast, presence of short-chain fatty acids may reduce the number of mosquito landings on an artificial bait [10]. Other studies explored the potential role of human blood type in host preference [11-13], but their findings remain conflicting and difficult to interpret – partly due to methodological challenges.

To this day it remains largely a mystery why some people receive more mosquito bites than others, even though bite frequency plays a key role in the risk of mosquito-borne diseases. Understanding factors behind mosquito attraction could improve prevention strategies in high-risk areas and possibly inform behavioural choices for those who would like to steer clear of mosquito bites. Existing cohort studies on mosquito attraction, however, are few, often use very small sample sizes, and show conflicting results, making them difficult to interpret. The present study, to our knowledge the largest of its kind, investigates why some people get bitten more than others in a large-scale ‘Mosquito Magnet Trial’ conducted during a 3-day music festival in the Netherlands. This setting was chosen because it pushes human behaviour, hygiene, and odour beyond the limits typically observed in observational studies.

## METHODS

### Study setting and participants

A Campingflight to Lowlands Paradise (or Lowlands) is an annual 3-day festival with 65,000 attendees that takes place on the farmlands of Biddinghuizen, The Netherlands. Our study was conducted in 2023 (August 18–20) in a custom-designed laboratory set up inside welded-together shipping containers. Whereas the festival for many festivalgoers very much is a 72-hour non-stop experience, participants were invited to join the study between 12 to 8 pm, when a science fair (Lowlands Science) was open for visitors. After being briefed about the study procedures, participants (≥ 18 years old) could voluntarily participate in the Mosquito Magnet Trial, which took around 20 minutes to complete. We consulted the national medical ethics review committee (Oost-METC) to assess whether ethical approval was necessary for this study. They issued a statement confirming that ethical approval was not required.

### Mosquito rearing and transport

Not unlike the festivalgoers, *Anopheles stephensi* mosquitoes [14] were reared on a reverse day-night rhythm under controlled conditions (at 30°C and 70-80% humidity with a 12-hour day/night cycle) at Radboudumc in Nijmegen, The Netherlands. Mosquito rearing occurred approximately 100 kilometres away from the study container at Lowlands festival in Biddinghuizen. Every morning during the study period, non-blood fed mosquitoes were transferred to netted cages, put in Styrofoam boxes, and transported by car to the study container in Biddinghuizen. Over the course of three days, a total of 1,700 mosquitoes were transported. At the study container and before the start of the study day (<12:00 midday), 20-35 mosquitoes were transferred to each of the study cages using an aspirator in a designated section of the container, separated from its surroundings by nets. For the remainder of the study day, mosquitoes remained in their cages.

### Study procedures and experimental design

Participants were asked questions about their general health, diet, and hygiene during the festival (hoping that this was different from their normal routine), and substance use (idem) in a questionnaire that was anonymous and offered in either Dutch or English. Subsequently, participants had their tympanic temperature taken and performed a breath alcohol test (Dräger Alcotest^®^ 7000, Zoetermeer, Netherlands). Participants then continued to the mosquito attraction test. To quantify mosquito attractiveness, we used a custom-designed video-based assay consisting of transparent acrylic mosquito cage (15 × 15 × 15 cm) housing 20-35 *Anopheles stephensi* mosquitoes [14] (Figure 1).

**Figure 1.**
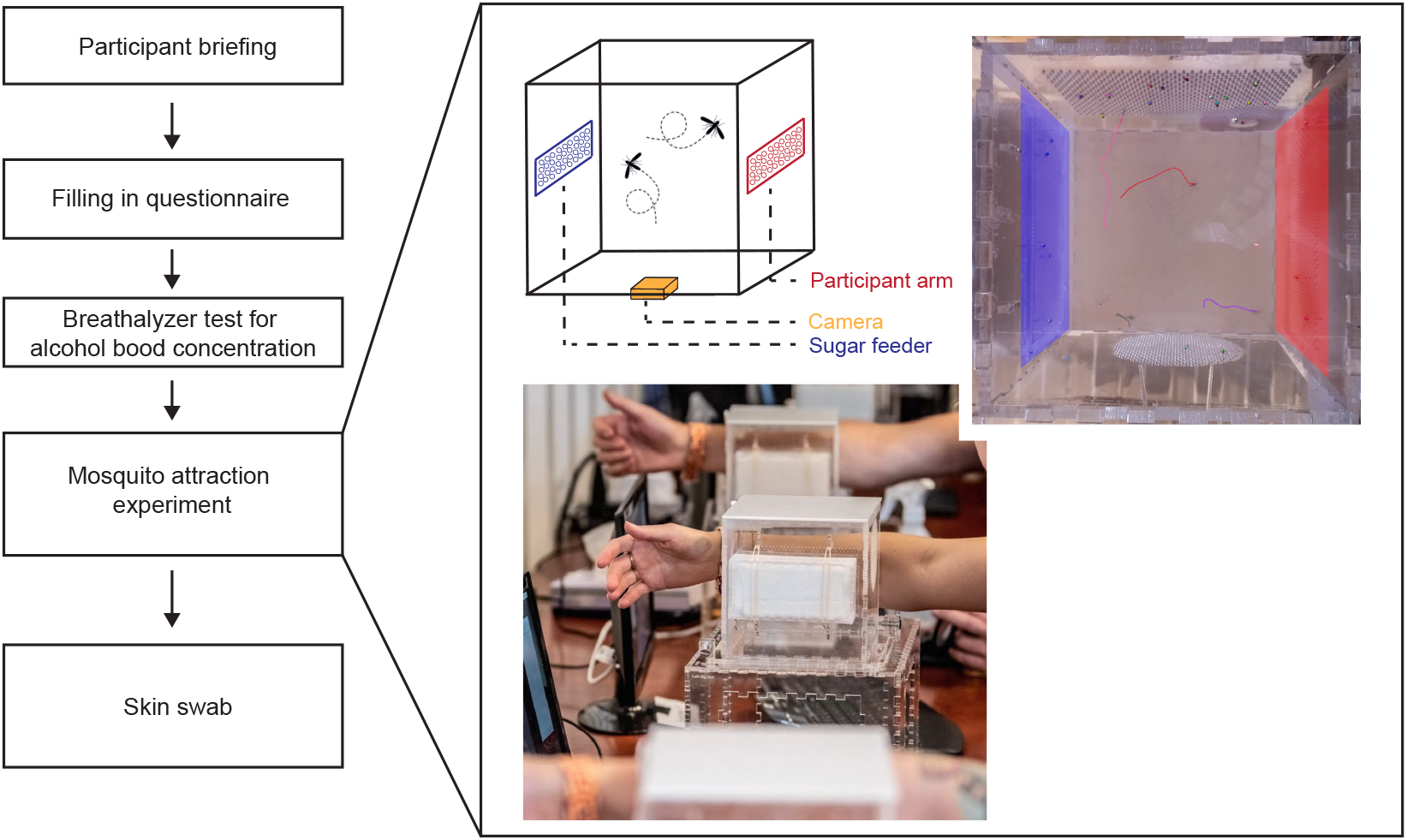
Schematic overview of study procedures. Participants were invited to fill in an anonymous questionnaire after being briefed about the study purpose and procedures. Alcohol blood concentration was measured using a Breathalyzer test. Mosquito attraction was quantified in a custom designed set-up consisting of an acrylic cubic cage housing 20-35 mosquitoes. Participants pressed their arms against the right side of the cage, where a grid of small perforations was located (highlighted in blue). Perforations were also located on the opposite side of the cage (the left side when facing the cage). Here, a sugar feeder was positioned (highlighted in red). A raspberry pi computer and associated camera were located beneath the transparent cage and recorded mosquito tracks for the duration of the experiment (3 minutes). Afterwards, participants were invited to perform a skin swab on their right forearm for skin microbiome assessments.

On two opposing sides, the cage had rectangular areas (12.5 × 5.5 cm) perforated with an array of holes (diameter: 0.8 mm, spacing: 3.2 mm). An acrylic box with cotton pads soaked with 10% glucose solution was attached to the left side of the cage. The mosquito cage was placed on a transparent acrylic base, equipped with an armrest and a Raspberry Pi microcomputer with associated camera (piCamera V2) situated beneath the cage to record flight and landing dynamics. Participants were seated at this setup and asked to gently blow into the cage for 10 seconds through perforations in the front panel. Next, participants were asked to present their lower right arm by holding it against the right perforated side of the cage, while placing their arm on an acrylic support to prevent motion and fatigue. Perforations in the cage wall allowed skin odorants to diffuse into the cage, while being sufficiently small to prevent mosquitoes from biting through. A three-minute video was recorded and stored for analysis. After use, mosquito cages were stored and reused after a resting period of approximately 20 minutes in Styrofoam boxes free from external stimuli.

### Video analysis and mosquito tracking

Using the pose estimation python package SLEAP [15], we fine-tuned a neural network to detect centroid coordinates of mosquitoes. We engineered a custom tracking algorithm based on a Kalman filter and the greedy tracking algorithm to track mosquitoes over time. Using the output of the tracking algorithm, we quantified the number of mosquitoes that are in flight or stationary at a given time. To do so, we employed a simple state machine that would take a track’s displacement between sequential frames and attribute the state of resting if the mosquito track did not change positions over a period of 7 frames, or attribute the state of flight to the mosquitoes that show displacement above a given threshold for more than 3 consecutive frames. With this data, not only did we quantify the number of resting and flying mosquitoes at a given point in time, we were also able to count the total number of landings on a given region of interest. On the 2D projection of the filmed cage, the left and right trapezoids represent each of our regions of interest (i.e. the area where the arm was presented and the sugar feeder area, Figure 1), while the remaining regions represent an out-of-region area (i.e. top and bottom trapezoids and center square). By using the position of these regions of interest, we quantified how many resting and flying mosquitoes are at each region of interest at any given time. Absolute attractiveness was quantified as the number of mosquito landings at the arm-region of interest, relative mosquito attractiveness was obtained by dividing the number of mosquito landings in the arm-region of interest by the number of landings on the sugar feeder region of interest.

### Skin swab

After quantifying mosquito attractiveness, participants received instructions on how to perform a skin swab. To prevent contamination, participants performed the skin swab themselves by rubbing a cotton-swab back and forth across an area of approximately 15 cm on the inner part of their left forearm. Participants were told to rub for approximately 30 seconds, after which they put the swab in a designated tube labelled with their anonymous participant number and filled with SCF-1 solution (50 mM TRIS, 1mM EDTA, 0.5% Tween 20). Tubes were transferred to zip-lock bags and stored in a −20°C freezer in the study container. After the festival, swabs were transported in Styrofoam boxes to the −80°C freezer at Radboudumc, Nijmegen.

### Laboratory assays and sample selection

To pilot 16S rRNA (16S) microbial sequencing using genetic material from skin swabs, we randomly selected 8 swabs from participants in the top two quartiles of relative mosquito attractiveness (Supplementary Figure 1). Microbial genomic DNA was extracted from swabs using the Zymo-BIOMICS DNA miniprep kit by vortexing the swab in a ZR BashingBead Lysis Tube with 750 µl of ZymoBIOMICS Lysis Solution for 30 seconds to bring the microbes into solution, followed by bead beating. Illumina 16S rRNA marker-gene (16S) amplicon libraries were generated and sequenced at BaseClear BV (Leiden, The Netherlands) on an Illumina MiSeq PE300 system. Sequencing of bacterial 16S rRNA is described in detail in the appendix. After confirming successful 16S sequencing in our pilot set, we selected an additional 85 swabs based on their relative mosquito attractiveness. This included 45 swabs from individuals with the highest daily attractiveness scores (15 per study day) and 40 swabs with the lowest daily attractiveness scores (12 from the first study day, and 14 from each of the next two days) after excluding samples with the daily lowest 10% of mosquito flight time and landings. The latter was done to avoid any bias from a very inactive cage resulting from overheating in the study container, or fatigued mosquitoes. The numbers used in this selection were arbitrary but avoided resource intensive analyses on skin swabs with a mediocre mosquito attraction score which were unlikely to inform our central question on whether skin microbiota differ between individuals who strongly attract mosquitoes and those who do not.

### Statistical analysis

The primary variable of interest was the number of arm-landings per mosquito. The relative attractiveness score (arm-landings divided by sugar feeding landings) was the secondary outcome variable of interest, and was used for the selection of skin swabs. We fitted generalized linear models with a log-link Gaussian distribution to investigate the relationship between arm-landings and a set of predictor variables divided in different categories. We excluded observations with a relative mosquito attraction score that fell in the daily bottom 10% of mosquito flight time and landing counts. This was done because, despite rotating the experimental cages to allow mosquitoes to rest, we observed a decline in mosquito responsiveness to participants’ arms over the course of the study day (Supplementary Figure 2). The number of arm landings per mosquito was set as the model’s outcome variable and a random effect with an interaction between time and day of the experiment was included to account for any potentially remaining temporal variation due to environmental temperature changes and mosquito fatigue. We additionally included an offset term with landings on the sugar feeder and anywhere else in the cage. This was done to model the effect on arm landings relative to the overall activity in the cage. Separate models were used to test fixed effects of variables categorized in i) demographics and health, ii) hygiene and care, iii) consumptions and diet, and iv) substance use. Participant age (in years) and time since last shower (in hours) were grouped into four quartiles and all categorical variables were included as factors. We selected fixed effects with a significant effect and included them in a final model to assess their combined effect. Since models shared the same outcome variable, we applied the Benjamini-Hochberg approach to adjust p-values for multiple testing (P_FDR_). Model coefficients were exponentiated to obtain fold changes that represent multiplicative changes in the outcome value. Mann Whitney U tests were used to test if continuous densities (e.g. number of consumed beers) differed between groups. A two-sample z-test for proportions was used to compare prevalences between groups (e.g. proportion of substance users). Alpha diversity metrics by the Shannon index were obtained from “qiime2 diversity core-metrics-phylogenetic” with default settings [16]. Principal Coordinate Analysis (PCoA) with weighted UniFrac distances was computed via “qiime2 diversity core-metrics-phylogenetic” [16] and the ‘vegan’ R package [17]. Permutational Multivariate Analysis of Variance (PERMANOVA) was applied on the weighted UniFrac distances to compute differences in skin microbiota beta diversity between selected study groups [18, 19]. The difference in relative taxon abundance between two study groups was tested with either the Mann-Whitney U test (non-paired) or Wilcox signed rank test (paired) with continuity correction in the normal approximation for the p-value. All statistical analyses were performed in R (version 4.4.1) and supported by substantial coffee consumption.

## Results

Despite being in steep competition with acts like Billie Eilish, Bombino, Underworld, and the North Netherlands Orchestra performing a riveting rendition of Beethoven’s 9th, our study attracted a total of 524 participants. Of these, seven individuals reported to have used mosquito repellents and were excluded from further analyses (Supplementary Figure 1). An additional 52 individuals were excluded because no video was recorded during the experiment (n = 17) or the mosquito activity fell in the daily bottom 10% of mosquito flight time and the daily bottom 10% of landing counts (n = 35).

Among the remaining 465 participants, a majority identified as female (55.7%, 259/465) compared to those who identified as male (43.0%, 200/465, P < 0.001, Figure 2). One individual reported to be non-binary, and five individuals did not indicate their gender in the questionnaire. A single participant identified as demigod; mosquitoes would not come near this participant and consequently she/he/they had to be excluded due to very low mosquito activity. Although the majority of individuals did not know their blood type (62.2%, 289/465), the most commonly noted blood type was type O (48.3%, 85/176). The proportion of males with blood type AB (14.1%, 9/64) was surprisingly high compared to the general national average of 3.0% [20]. Blood type was self-reported and not confirmed through laboratory testing, it thus may be subject to inaccuracies in participants’ knowledge or recall. Sunscreen usage was high with over half of participants indicating to have used sunscreen on their forearm that day (52.7%, 244/463). The majority of participants did not have a special diet (77.2%, 338/438), followed by vegetarians (13.9%, 61/438), and pescatarians (5.7%, 25/438). Whether out of genuine love or mere convenience, canned sausages stole the culinary spotlight, appearing in 15.6% of people’s latest meals (36/231). Participants used a wide variety of substances, with cannabis being the most frequently reported (19.6%, 91/465) followed by XTC (18.7%, 87/465) and cocaine (11.4%, 53/465) and no difference between males and females regardless of substance type (all P > 0.279). A non-negligible proportion of participants reported use of multiple substances with two reporting all listed substances except for amphetamines in the past 48 hours. The majority of participants drank at least one beer in the past 12 hours (65.6%, 299/456), this proportion was even higher amongst men (79.8%, 158/198). A higher proportion of beer abstainers reported sleeping alone last night (61.1%, 96/157) compared to beer drinkers (41.4%, 103/249, P < 0.001), yet all four individuals who consumed the most beers (> 20 beers in the past 12 hours) also slept alone. While beer consumption may not guarantee good decisions, it might at least facilitate company when consumed in moderation.

**Figure 2.**
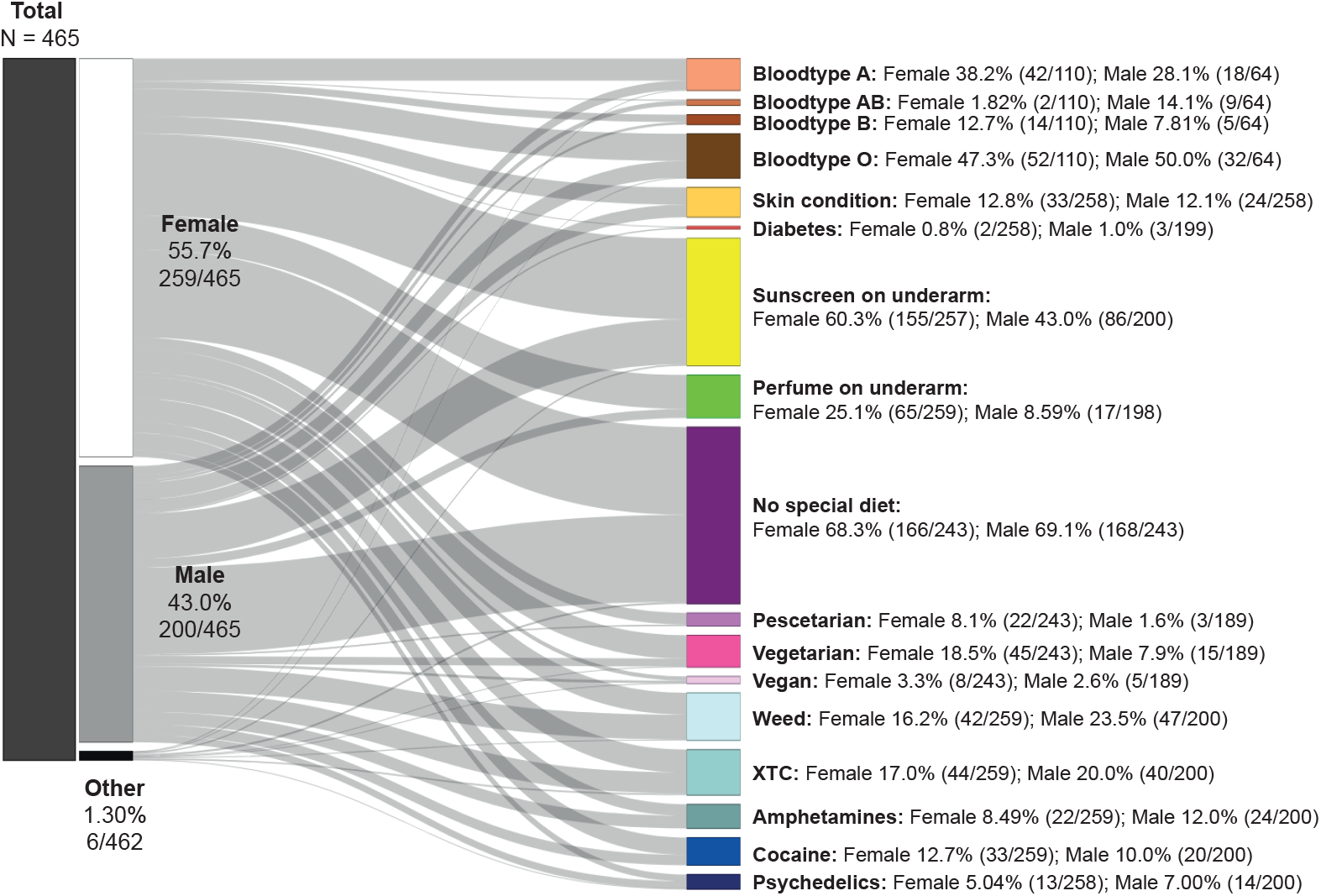
General cohort characteristics. Sankey plot showing the proportion of participants grouped into different categories. Bar sizes reflect the total proportion, whereas the proportion per participant sex (female/male) is indicted in text.

Mosquito attraction varied across participants, with the number of arm-landings per mosquito ranging from 0 to 14.8 (Supplementary Figure 3). Surprisingly, three of the four individuals with zero arm-landings came to the study container during the performance of Joost Klein, who was then famous for his song ‘*Friesenjung*’, a hardcore take on a parody of Sting’s *‘Englishman in New York’*. Contemplating on whether mosquitoes could be repelled by upbeat electronic music, we continued assessing factors behind mosquito attraction using multivariate linear models (Figure 3). Mosquito attraction was similar between female and male participants (P_FDR_ = 0.189, Figure 3A). We found evidence for an interaction between time since last shower (in quartiles) and sunscreen on a participant’s forearm (all P_FDR_ < 0.007). Amongst participants that showered recently (<6 hours ago), the presence of sunscreen on the forearm made them less attractive (Fold change [FC] 0.52, 95% CI 0.38-0.71, P_FDR_ < 0.001, Figure 3B). The repellent effect of sunscreen became less pronounced when time since last shower increased. We found no effect of sunscreen usage when applied anywhere but on their forearms (P_FDR_ = 0.992). We did not observe an effect of perfume when applied on participant’s forearm or elsewhere (both P_FDR_ > 0.641), potentially because of abundant olfactory triggers. Interestingly, participants who attracted a partner into their tent the night before also attracted more mosquitoes (FC 1.46, 95% CI 1.24-1.71, P_FDR_ < 0.001), without evidence for an interaction with time since last shower (all P_FDR_ > 0.182).

**Figure 3.**
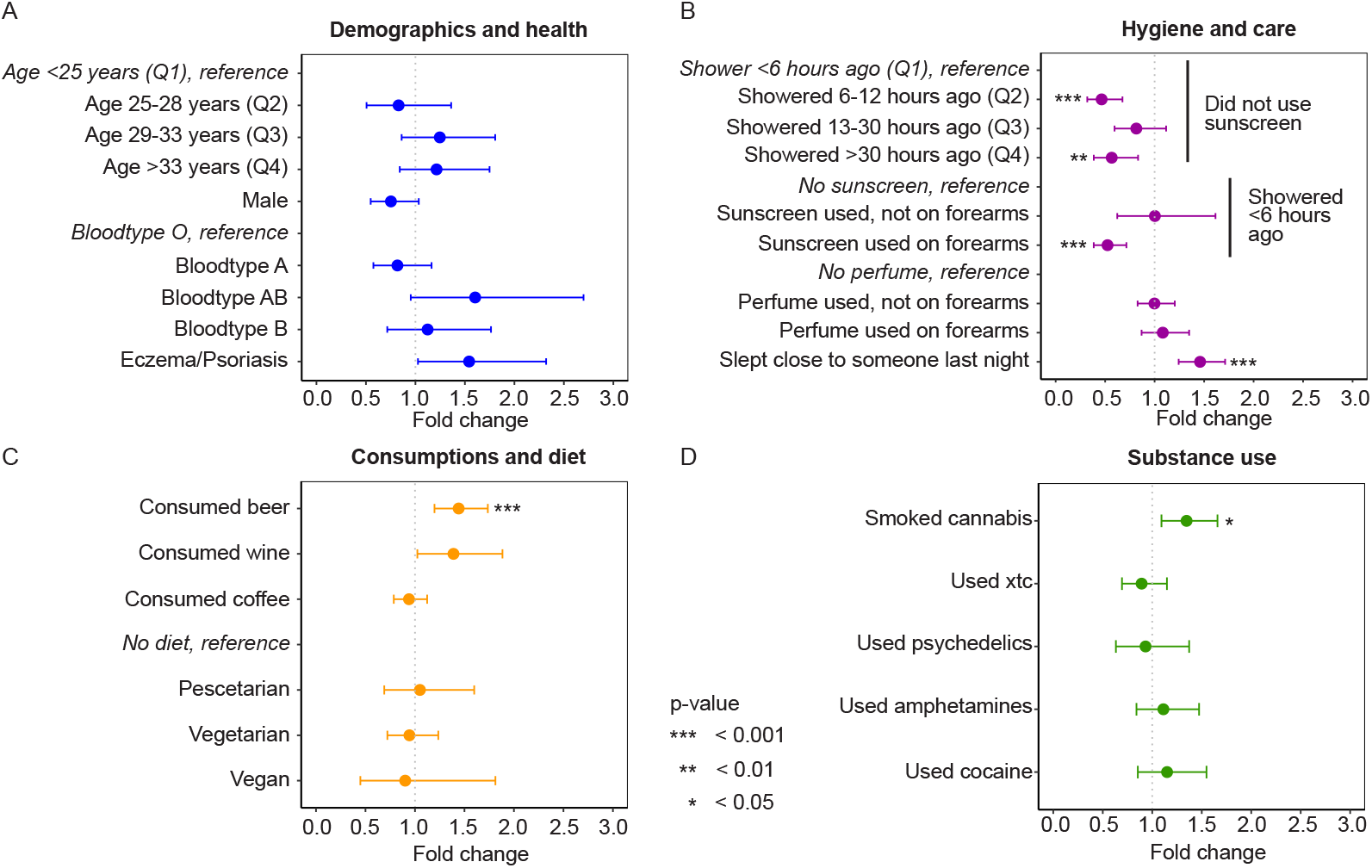
Category-wise assessment of factors influencing mosquito attraction. Forest plots showing fold changes in mosquito attraction (arm landings) for different participant characteristics relative to reference groups. Variables were grouped into different categories (A-D). A fold change greater than one indicates an increased attraction compared to reference. Dots represent the point estimates, horizontal bars indicate the 95% confidence intervals (CI). Asterisks represent significance levels of <0.05 (*), <0.01 (**), and < 0.001 (***), after adjusting for multiple testing.

Mosquitoes showed a clear preference for the well-hydrated, on hops and grapes, that is. Arm landings were significantly higher in beer drinkers compared to those who had nobly abstained for at least 12 hours (FC 1.44, 95% CI 1.20-1.74, P_FDR_ < 0.001, Figure 3C). Mosquitoes seemed to have a taste for wine drinkers too (FC 1.39, 95% CI 1.02-1.88, P = 0.035), but this effect sobered up after correcting for multiple testing (P_FDR_ = 0.103). Measured blood alcohol concentration ranged from to 1.82‰ and positively correlated with the self-reported consumed number of beers (Spearman rho = 0.46, P < 0.001) and glasses of wine (Spearman rho = 0.12, P = 0.011). No statistically significant effect of alcohol concentration was observed on mosquito attraction when included as a continuous variable (FC 1.04, P_FDR_ = 0.853) nor as a binned variable using the concentration of approximately two units as a threshold (< 0.5‰ versus 0.5‰, FC 1.21, P_FDR_ = 0.344). Individuals reported to have smoked cannabis in the past 48 hours were more attractive to mosquitoes than individuals that did not smoke cannabis (FC 1.35, 95% CI 1.09-1.66, P_FDR_ = 0.017, Figure 3D). Cannabis was the only substance for which an effect on mosquito arm-landings was found, the effects of other substances were statistically not significant (all P_FDR_ > 0.569). There was no indication that the presence of a cannabis user made mosquitoes fly at higher altitudes or made them less aggressive.

We then assessed the effect of statistically significant variables from different categories combined in a single model. Mosquito arm-landings were still higher for individuals that drank at least one beer in the past 12 hours (FC 1.35, 95% 1.12-1.63, P_FDR_ < 0.001, Figure 4); the effect of cannabis remained positive (FC 1.18, 95% 0.97-1.44) but lost statistical significance (P_FDR_ = 0.216). Individuals who had slept with someone else the night before were 1.34 times more attractive than individuals who slept without close company (95% CI 1.14-1.58, P_FDR_ = 0.002), without evidence for an interaction with beer consumption (P = 0.971). The interaction between time since latest shower and application of sunscreen on forearms remained robust (P_FDR_ < 0.001). Individuals that showered recently and used sunscreen on their forearms were still less attractive than individuals who did not use any sunscreen (FC 0.52, 95% 0.38-0.70, P_FDR_ < 0.001).

**Figure 4.**
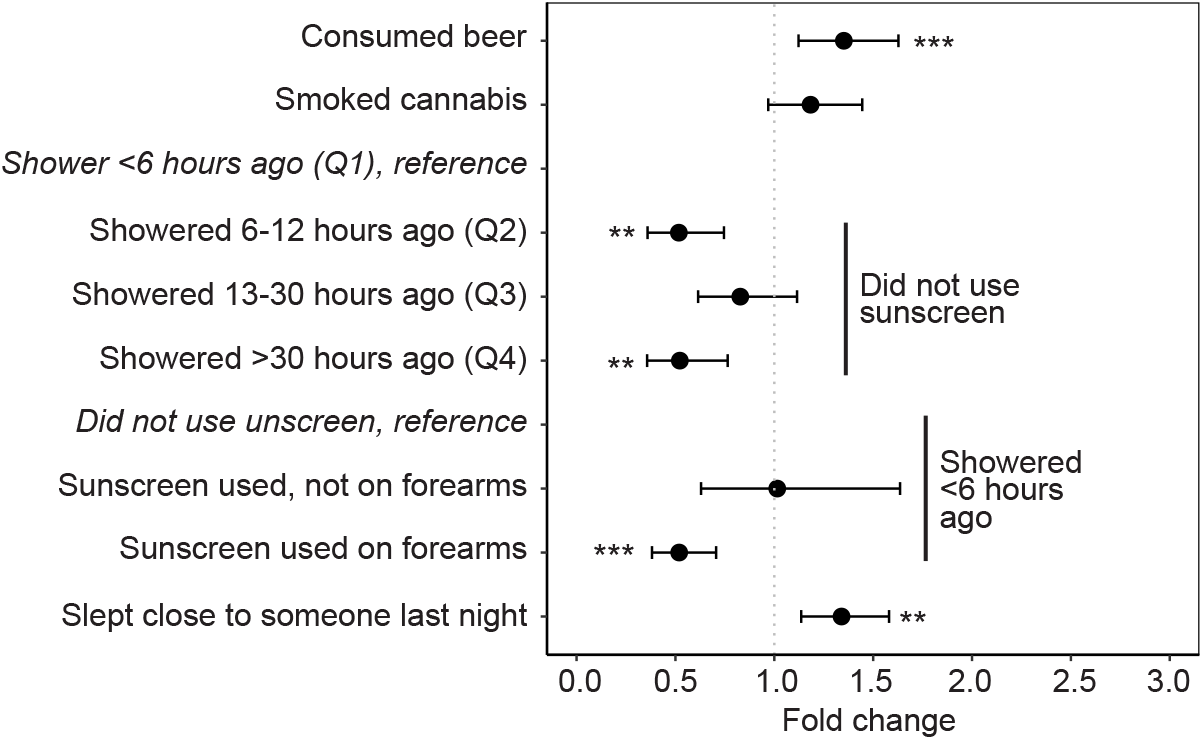
Factors driving mosquito attraction. Statistically significant variables (corrected for multiple testing) were selected from the category-wise models and assessed in a single model. Forest plot showing fold changes in mosquito attraction (arm-landings) for different participant characteristics relative to reference groups. A fold change greater than one indicates an increased attraction compared to reference. Dots represent the point estimates, horizontal bars indicate 95% confidence intervals (CI). Asterisks represent significance levels of <0.05 (*), <0.01 (**), and < 0.001 (***), after adjusting for multiple testing.

A variety of bacterial taxa were detected on forearm skin swabs from selected individuals (Supplementary Figure 4). The alpha diversity of the skin microbiota, indicated by the Shannon index, did not differ between the subset of highest and lowest attractive individuals (P = 0.77, Figure 5A). We also found no evidence of a different overall microbiota composition based on beta diversity between high and low attractive individuals (PERMANOVA P = 0.232, Supplementary Figure 5). The four most dominant genera in both high and low attractive individuals were *Cutibacterium, Sphingomonas, Corynebacterium*, and *Staphylococcus* (Figure 5B). All of these genera are recognized as skin commensals except for *Sphingomonas*, which is associated with water and other environmental sources [21, 22]. Its presence on the skin may indicate environmental exposure. The dominant presence of *Corynebacterium*, previously linked to human malodour [23, 24], suggests a distinctive smell in our study container, though we will refrain from evaluating individual contributions. *Streptococcus* was found to be more abundant on high compared to low mosquito-attractive individuals (2.68% vs. 0.61% mean relative abundance, respectively), although the effect was modest and only statistically significant without correction for multiple testing (Mann Whitney U P-uncorrected = 0.017).

**Figure 5.**
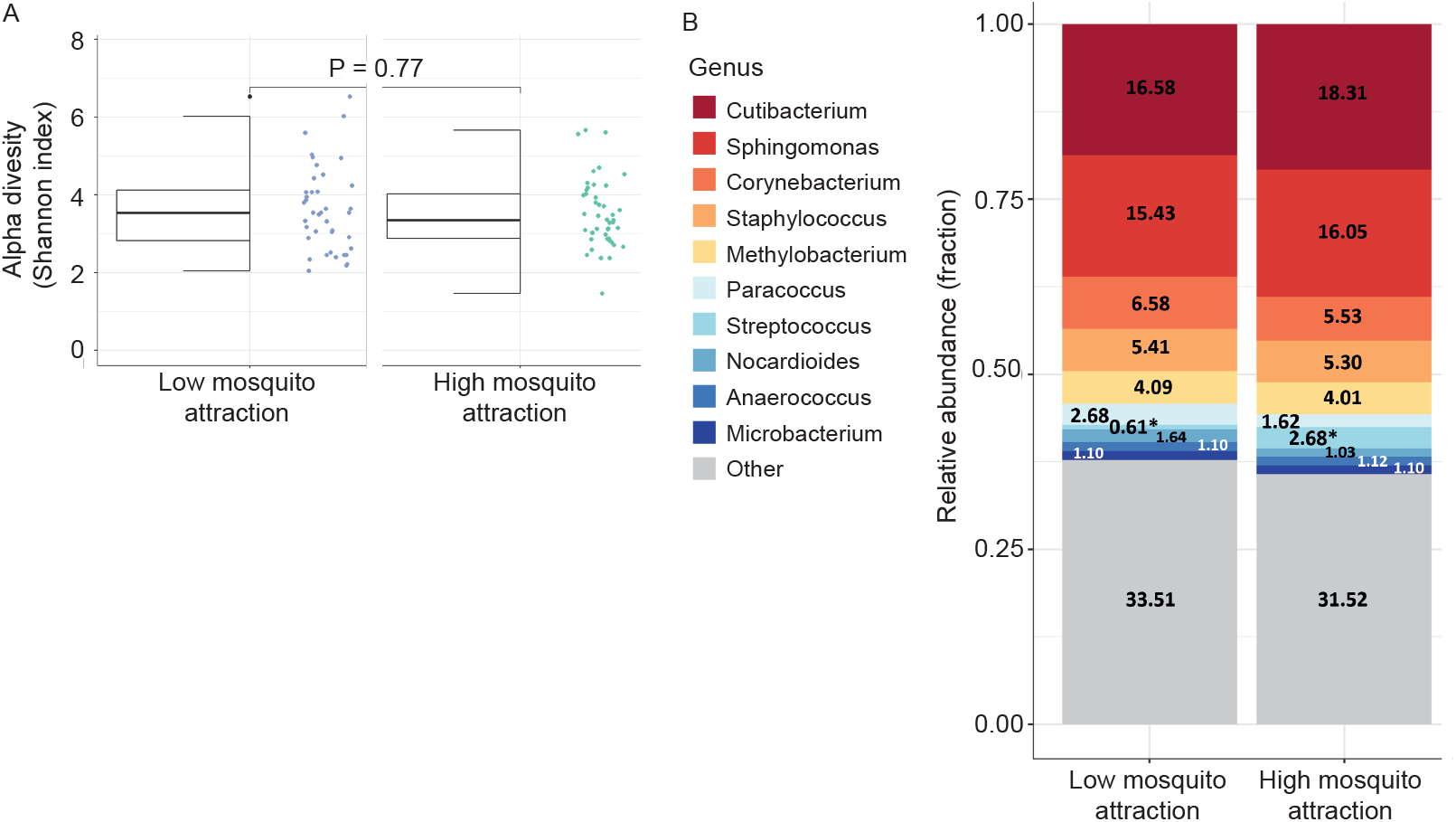
Forearm skin microbiome diversity and composition in relation to mosquito attractiveness. A subset of 85 participants with highest (n=45) and lowest (n=40) mosquito attractiveness were selected for skin microbiome assessments using forearm skin-swabs. A) Alpha diversity box plots of study groups with lowest (left) and highest (right) relative attraction scores. Alpha diversity was assessed using the Shannon index and no significant difference in diversity was observed. B) Stacked-bar graphs with the average genus-level relative abundance values for low (left) and high (right) attraction score individuals. Shown are the top 10 most abundant genera for low versus high attractors. The average successful classification rate up-to genus level was 88.6% ± 6.1% SD, for the complete dataset of n=97 samples (including 8 pilot samples).

## Discussion

Some people are bitten by mosquitoes more often than others, but the underlying mechanisms remain poorly understood. In the present study, we developed an experimental setup to investigate differences in mosquito attraction in a cohort of festival goers in the Netherlands. Mosquito attraction was quantified by the number of landings on the participant’s arm-side of the experimental cage, revealing substantial heterogeneity across the cohort.

Beer consumption significantly increased mosquito attraction. This observation is consistent with previous work studying mosquito attraction in a different experimental set-up with smaller sample sizes [25, 26] and a considerably more modest intake. Measured blood alcohol concentration did not have an observed effect on mosquito attraction. Mosquito may simply be drawn to the distinctive smell of a Heineken pilsner, and alcohol blood concentrations may also be high because of wine or liquor consumption. Given the number of beer drinkers in our study, it’s clear that beer holds a special place for many. That said, while our findings are intriguing, there may be other contributing factors yet to be identified. Further research is certainly needed before we suggest rejecting a beer solely to avoid mosquito bites.

We also observed a negative association between the application of sunscreen on participants forearm and mosquito attraction. This association became less negative when participants showered a longer time ago. We did not ask participants when they applied sunscreen, but we assume it would be after their latest shower. The observed interaction suggests that sunscreen’s negative effect on mosquito attraction may wane over time. The observed effect could stem from sunscreen blocking human odours naturally present on a participant’s arm, or the sunscreen could contain a mosquito repellent component. This component is likely generally present in different sunscreen types and brands, given the consistently lower mosquito attraction observed and the likelihood that participants used a variety of sunscreen products. Even though further studies are needed to confirm and elaborate on the effect of sunscreen, it may be wise to bring some to the next summer barbeque.

We did observe a modest difference in skin microbiota of individuals with high versus low mosquito-attractiveness. In line with a previous report [27], we observed *Streptococci* to be significantly more abundant in highly attractive individuals when uncorrected for multiple testing. Although using different set-ups like sampling feet instead of arms and excluding alcohol intoxication, earlier work found elevated levels of *Staphylococci* in the skin microbiome of individuals highly attractive to mosquitoes [6, 27]. One of these studies did report an additional difference in bacterial diversity [6], whereas this was not confirmed by the other [27] and neither by our study, highlighting the complex role of the skin microbiome in human odour profiles and associated mosquito attraction. Importantly, others applying 16S rRNA sequencing to identify skin microbes have done this in a controlled lab setting where participants were, for instance, not allowed to shower on the day of sampling or use a sauna within 48 hours beforehand [28]. Our natural study setting with a ‘pop-up’ laboratory container situated in a steamy farmland populated by sixty thousand festival attendees limited control over these variables. Our participants were allowed to have showered, a privilege some clearly declined, and the music festival even had a sauna that people could have gone to.

Our study has several limitations. Although the large scale of our study distinguishes it from previous work, its setup also markedly differs from the controlled conditions used in earlier mosquito attraction studies [6, 9, 13, 25-27], thereby limiting direct comparison of findings. Admittedly, though we like to think there’s a curious inner scientist in everyone, our sample of science-loving festival-goers is not likely to be the average cross-section of humanity. Moreover, whilst substance use and alcohol uptake was already high in our cohort, it may be underestimated compared to the general festival crowd, considering the most active and perhaps least responsible partygoers (i.e. those frequenting the 24-hour stage) could still be asleep or too busy partying during our data collection hours.

The general picture that emerges from our study suggests that a sober life-style — abstaining from drugs and alcohol, sleeping alone, and applying sunscreen regularly — lowers one’s chances of getting bitten by mosquitoes. While we found no evidence supporting popular myths such as blood type influencing bite frequency, we were unable to assess the existence of so-called ‘sweet blood.’ Ultimately, enjoy the next festival or camping trip as you like—but it seems mosquitoes may have a soft spot for those making less responsible choices.

## Acknowledgements

We extend our gratitude to all individuals involved in the Lowlands Mosquito Magnet Trial logistics. We would like to thank Jolanda Klaassen, Laura Pelser-Posthumus, and Astrid Pouwelsen for breeding of the mosquitoes at Radboudumc. We also thank Wouter Graumans, Laura Akkerman, and Matthijs Jore for their assistance with mosquito transport to and from the study site. We are also thankful for the supply of Breathalyzer tests (Dräger Alcotest^®^ 7000), kindly provided by Dräger Nederland B.V. (Zoetermeer, Netherlands). We thank the Lowlands Science crew for giving us the opportunity to perform the study at their festival, and highly appreciate the support of Roos Rodenburg and others in facilitating the study container setup. Finally, we are deeply grateful to all participants who took part in our study and to those who excitingly shared their perspectives on being mosquito magnets. This work was supported by a VIDI (grant VI.Vidi.213.167) and XS grant (OCENW.XS23.4.114) from NWO (the Dutch science foundation)

All authors declared to have no competing interests according to ICMJE uniform disclosure forms.

---

**Supplementary Figure 1:** Sample selection procedures. The number of participants included at start, after exclusion due to i) absence of video records, ii) low mosquito activity, or iii) usage of mosquito repellent, and those selected for skin-swab assessments.

**Supplementary Figure 2:** Density plots indicating the number of landings per mosquito outside any region of interest, landings on the sugar feeder and landings on the participants arm region, separated by time of the experiment (in slots of 2 hours).

**Supplementary Figure 3:** Density plot showing the number of mosquito arm-landings across all included participants. The number of arm-landings per mosquito ranged from 0 to 14.8 in one measurement of three minutes.

**Supplementary Figure 4:** The overall microbiota composition of forearm skin samples. Sankey visualisation of the overall 16S microbiota composition of selected forearm skin samples (n=93). Values represent taxa average relative abundances over all samples, at different phylogenetic levels from phylum (left) up-to genus level (right).

**Supplementary Figure 5:** 16S analysis of the forearm skin microbiota beta diversity. Principal Coordinate Analysis (PCoA) of the Beta diversity based on weighted UniFrac distance, on study groups with low (red) and high (cyan) attraction score individuals (PERMANOVA p-value = 0.232).

