## Supplementary figures and images for "Blood, sweat, and beers: investigating mosquito biting preferences amidst noise and intoxication in a cross-sectional cohort study at a large music festival"

### Supplementary Figure 1

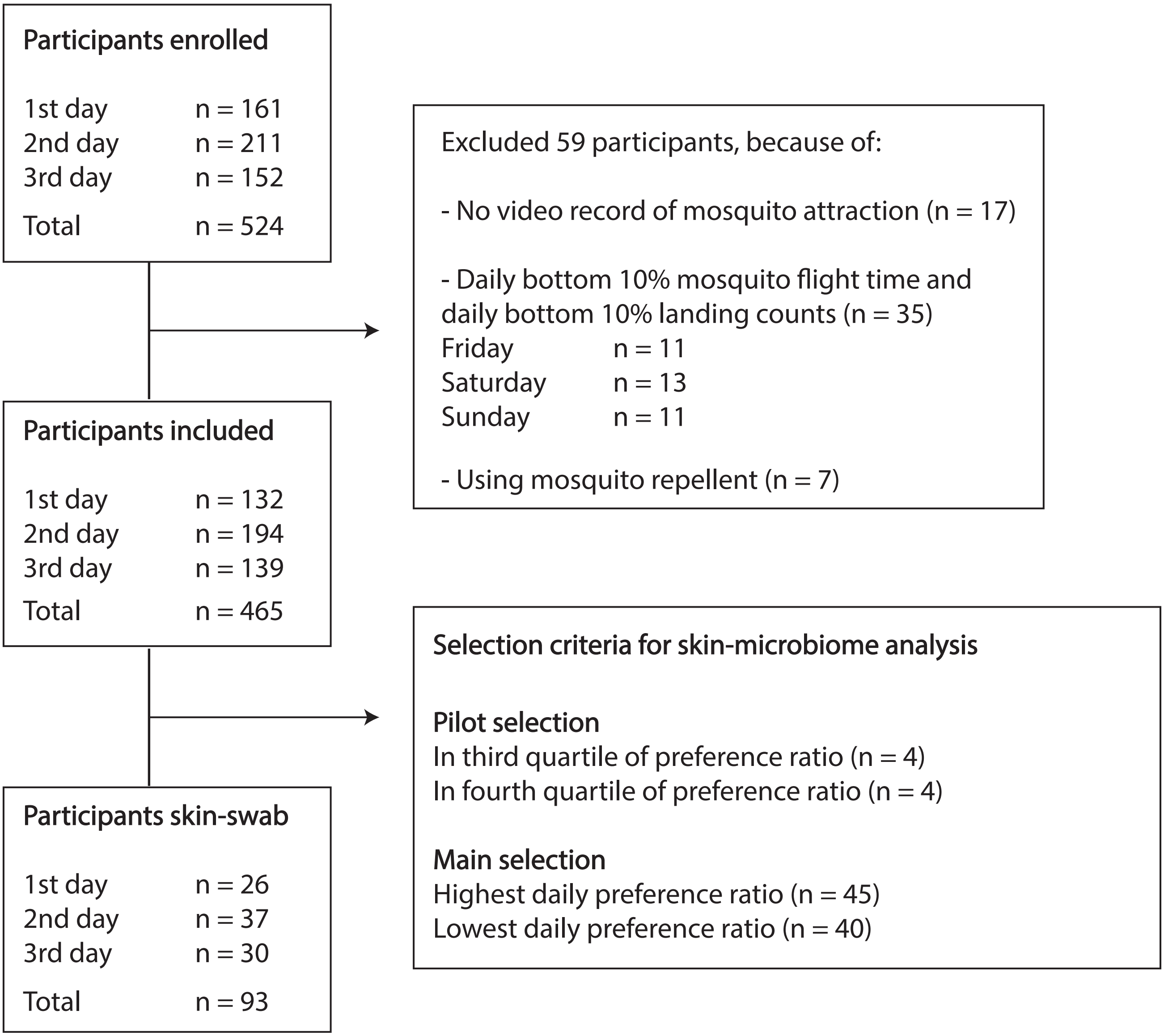

### Supplementary Figure 2

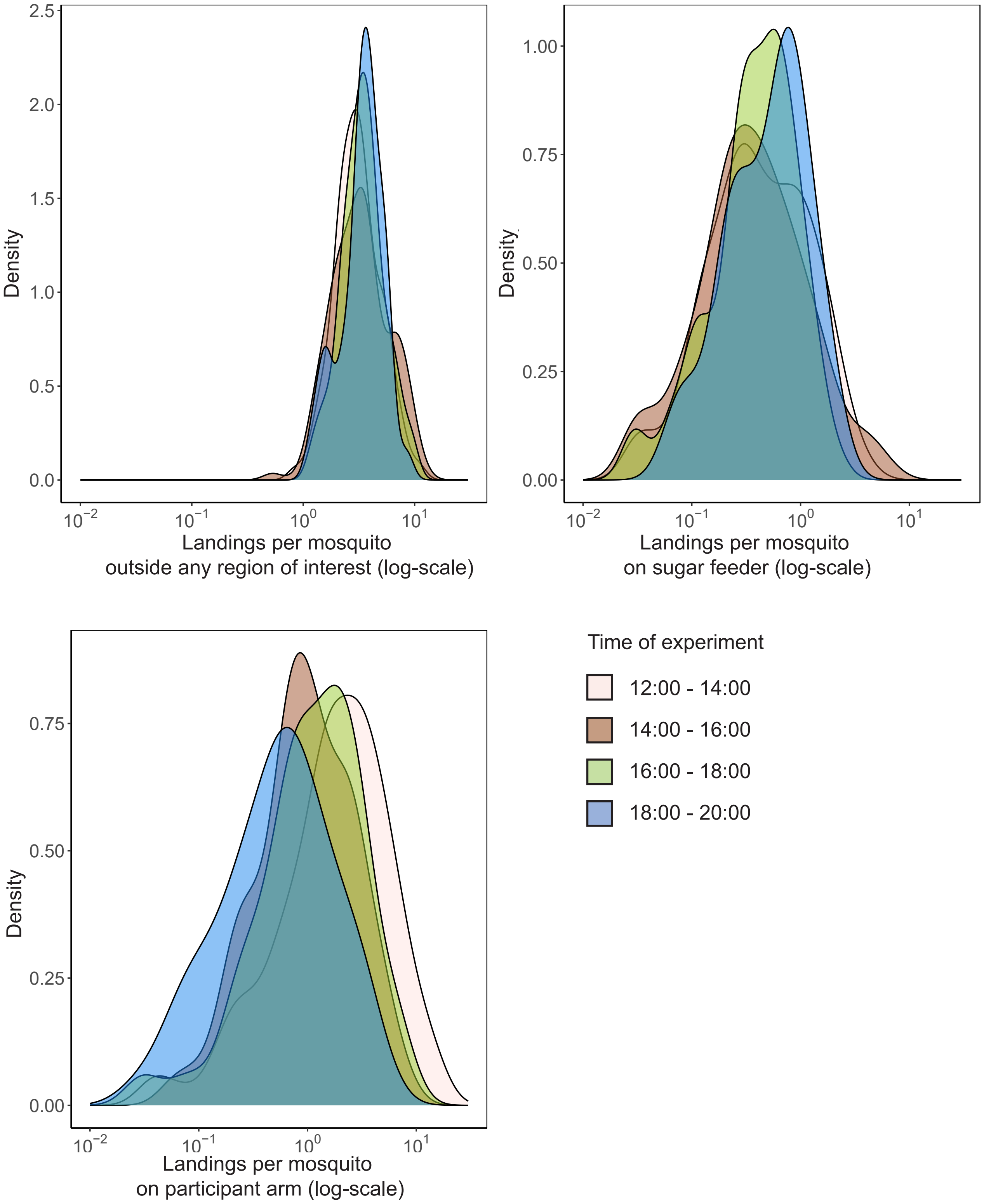

### Supplementary Figure 3

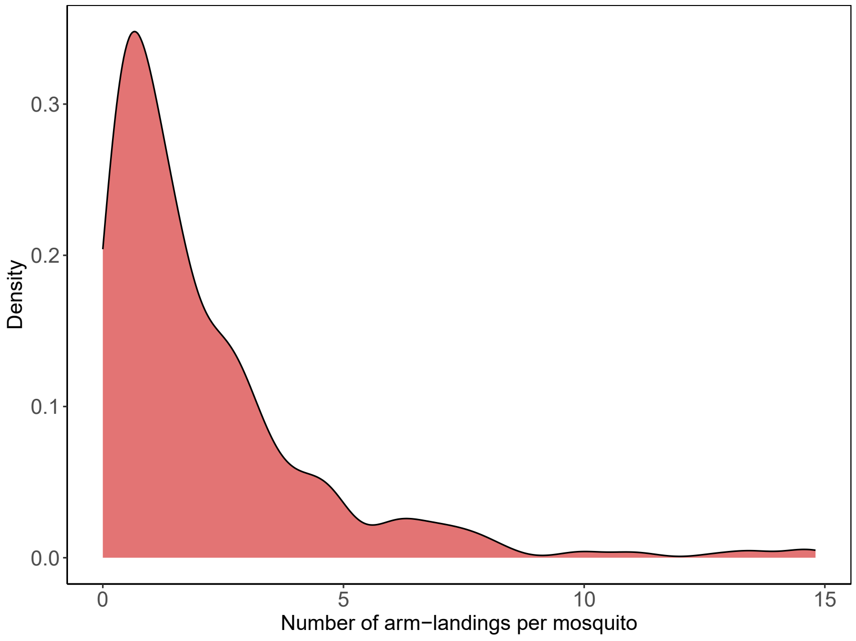

### Supplementary Figure 4

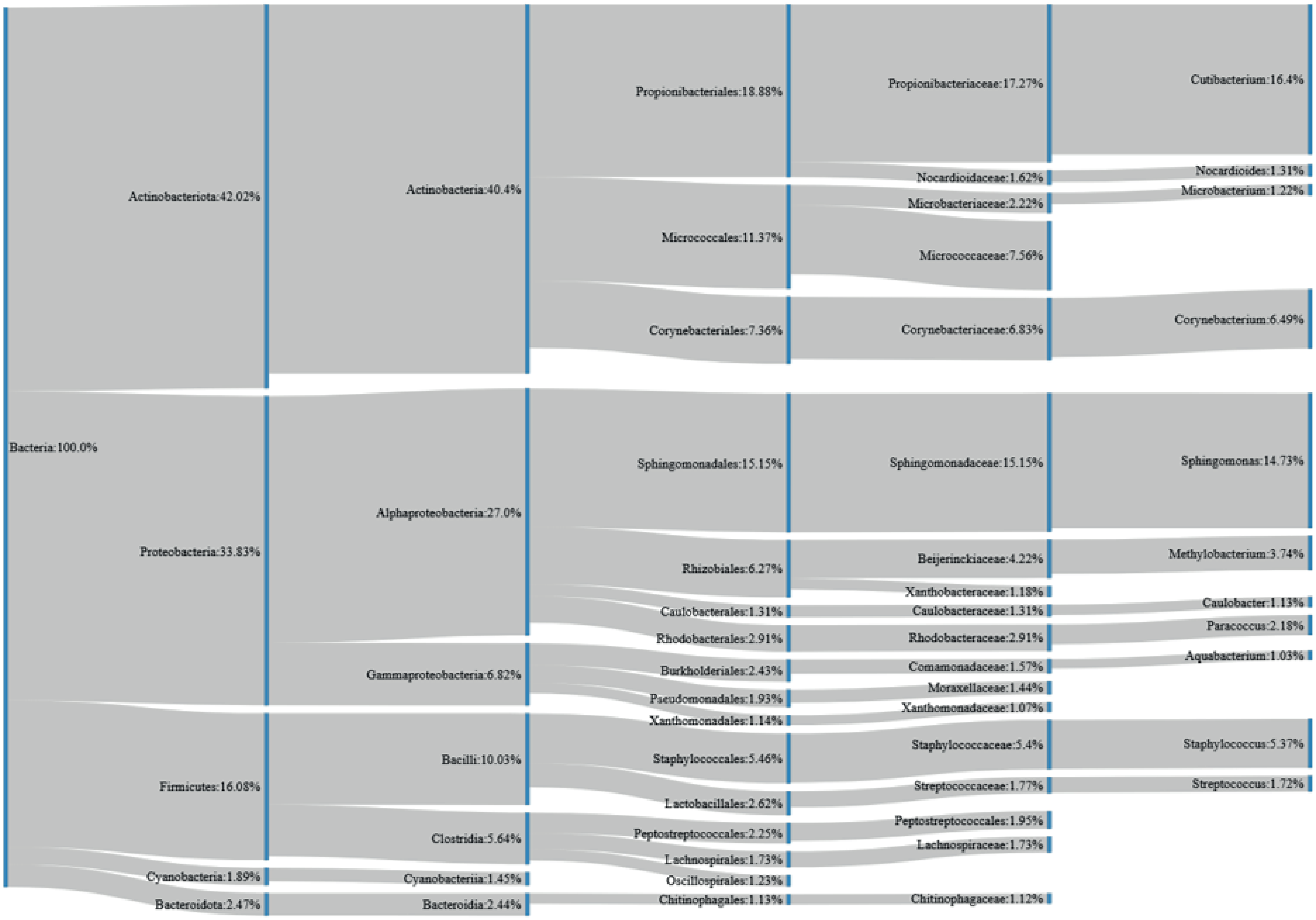

### Supplementary Figure 5

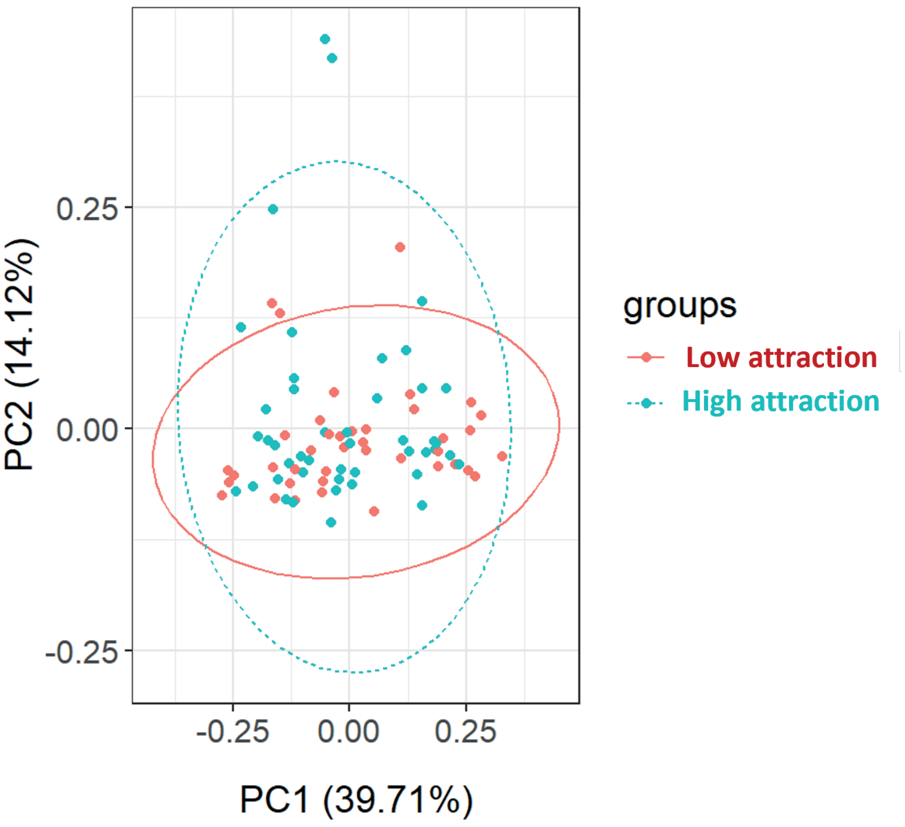
